# Network pharmacology, bioinformatics, molecular docking and experimental verification of the mechanism of HXZTS treatment on osteoarthritis synovium

**DOI:** 10.1101/2025.09.25.678423

**Authors:** Li Chen, Hongxiu Wang, Dongzhi Wu, Jinlan Su, Shunxi Chen, Dongdong Chen, Tao Zhang, Wenhui He

**Author notes:** Correspondence: (Tao Zhang); (Wenhui He). These authors contributed equally to this work and share the first authorship (Li Chen, Hongxiu Wang).

## Abstract

**Background:** Osteoarthritis (OA) is a prevalent joint disease characterized by synovial inflammation and synovial fibroblast dysfunction. Elucidating the molecular mechanisms underlying synovial tissue impairment in OA is crucial for developing targeted therapeutic interventions. Huoxuezhitongsan (HXZTS), a traditional Chinese medicine formula, has shown promising therapeutic effects in treating OA, but its mechanisms of action on synovial tissue remain unclear.

**Method:** This study integrated network pharmacology, bioinformatics analysis, molecular docking, and experimental validation to elucidate the potential targets, pathways, and mechanisms by which HXZTS exert therapeutic effects on osteoarthritic synovial tissue.

**Results:** Network analysis identified 21 active ingredients and 111 common targets of HXZTS for OA, with key compounds like quercetin, β-sitosterol, and stigmasterol. Enrichment analysis revealed HXZTS modulates inflammation through TNF, IL-17, and MAPK signaling pathways, oxidative stress, apoptosis, angiogenesis, and metabolic regulation. Molecular docking demonstrated strong binding affinities of HXZTS compounds with targets like AKT1, TNF, TP53, IL6, and VEGFA. RT-qPCR validation confirmed the downregulation of inflammatory mediators like IL6 and PTGS2, alongside the upregulation of genes associated with potential cartilage repair effects, such as MMP1 and MMP3, upon HXZTS treatment.

**Conclusions:** HXZTS exerts therapeutic effects on osteoarthritic synovial tissue through a multifaceted mechanism involving modulation of inflammation, oxidative stress, apoptosis, angiogenesis, and metabolic regulation. These findings provide insights into the potential application of HXZTS as a complementary or alternative therapeutic approach for managing OA and a foundation for developing novel targeted therapies.

## 1. Introduction

Osteoarthritis (OA) is a prevalent joint disease affecting approximately 528 million people worldwide [1]. OA is characterized by symptoms such as joint pain, stiffness, and swelling, which pose a significant health burden globally [2]. Current treatment strategies primarily focus on symptom control and improved joint function, ranging from lifestyle changes and physical therapy to pharmacological interventions, with joint replacement surgery for severe cases [3]. Therefore, early diagnosis and timely prevention of OA hold significant practical relevance. However, the pathogenic mechanisms underlying OA remain unclear. While past research has largely centered on articular cartilage, emerging evidence implicates the synovial membrane, synovial fibroblasts, and subchondral bone cells in OA pathogenesis as well [4]. The synovial membrane, a thin tissue lining the joint cavity, plays a crucial role in joint health and disease [5]. Synovial inflammation and changes in synovial fibroblasts have been linked to OA-associated joint degeneration and pain [6, 7]. Understanding the molecular mechanisms underlying functional impairment of the OA synovial tissue is critically important for developing targeted therapeutic interventions that can effectively modify disease progression and alleviate symptoms.

Traditional Chinese medicine (TCM) has been used extensively for thousands of years in China to treat inflammatory diseases. Huoxuezhitongsan (HXZTS) is a classic TCM formula composed of six herbs: Danggui (*Angelica sinensis* (Oliv.) Diels, DG), Sanqi (*Panax notoginseng* (Burkill) F.H.Chen, SQ), Ruxiang (*Boswellia sacra Flück.*, RX), Bingpian ( *Cinnamomum camphora* (L.) Presl, BP), Tubiechong (Ground Beetle, TBC), Zirantong (Pyrite, ZRT) (DG, SQ, RX and BP names have been checked with MPNS (http://mpns.kew.org)). The traditional function of HXZTS is to promote blood circulation, remove blood stasis, reduce swelling, and alleviate pain. HXZTS has been widely used in traditional Chinese medicine for the treatment of trauma-related pain and blood stasis-related disorders. In recent years, Qinbo et al. observed significant therapeutic effects of HXZTS in the treatment of knee osteoarthritis [8]. However, the molecular mechanisms underlying its therapeutic effects on OA synovial tissue remain unclear.

With the rapid development of systems biology and bioinformatics, network pharmacology has emerged as a powerful tool for studying the mechanisms of Chinese medicine formulas [9]. By integrating oral bioavailability prediction, drug targeting, and enrichment analysis, network pharmacology can identify active compounds and predict drug-target interactions and biological pathways [10]. Recently, several studies have applied network pharmacology to explore the mechanisms of Chinese medicine formulas for treating rheumatoid arthritis and osteoarthritis [11, 12]. However, network analysis of HXZTS on OA synovial tissue has not been reported yet.

Therefore, in this study, we aim to elucidate the pharmacological mechanism of HXZTS on OA synovial tissue through the integration of network pharmacology and bioinformatics methods, molecular docking, and experimental validation. The flow chart of the entire study is shown in Fig. 1. We explored the active ingredients, potential targets, and signaling pathways of HXZTS on OA synovial tissue, and verified it through molecular docking and experiments. It may also offer novel therapeutic targets for the discovery and management of osteoarthritis and related inflammatory diseases.

**Fig. 1.**
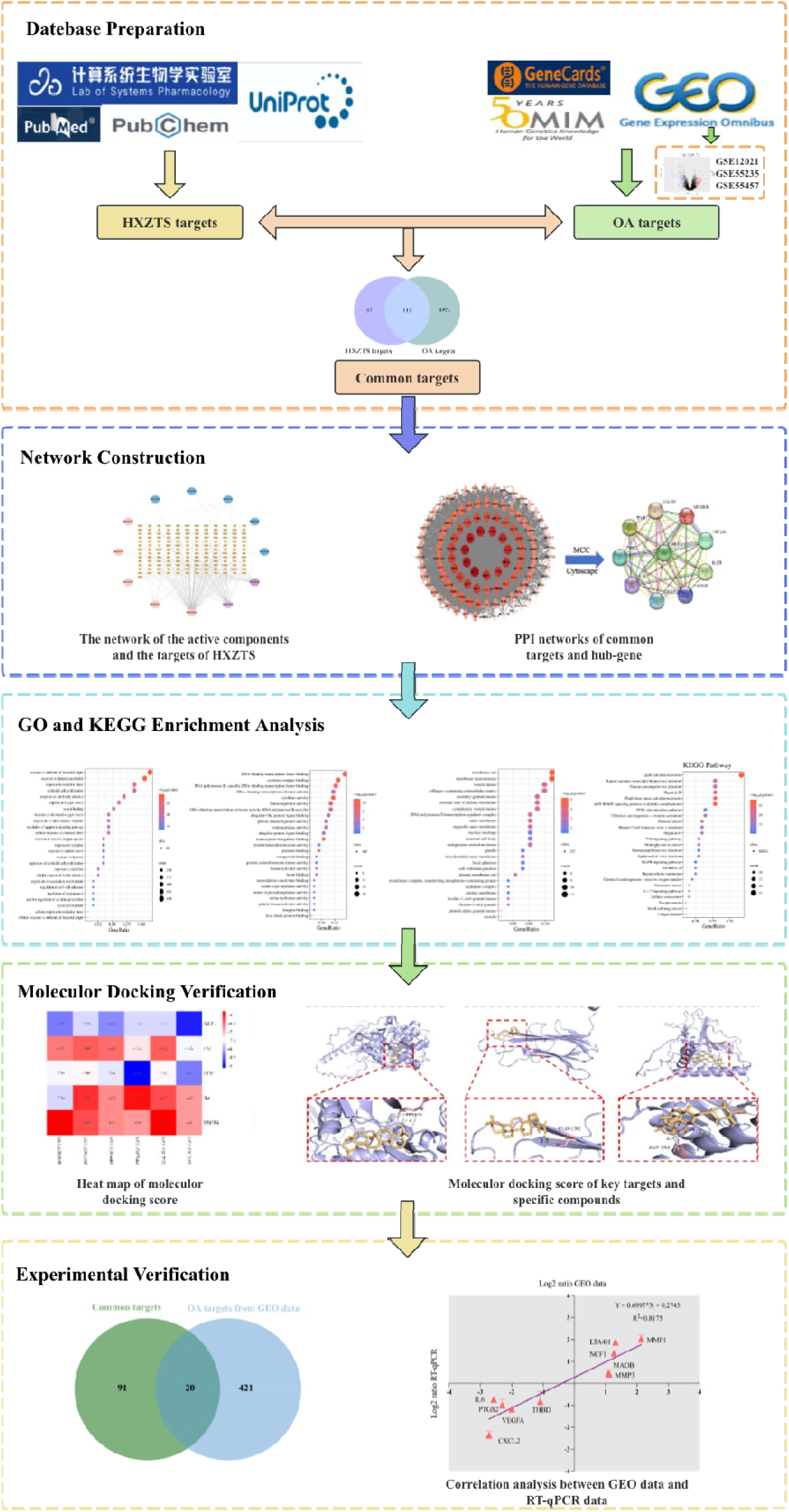
**The graphical abstract of exploring the mechanism of HXZTS in treating osteoarthritis synovitis.**

## 2. Materials and methods

### 2.1 Data collection

#### 2.1.1. Screening of active compounds of HXZTS

Data for DG, SQ, RX, and BP were obtained from the TCMSP database (https://old.tcmsp-e.com/tcmsp.php) [13]; data for TBC and ZRT (whose main component is ferrous disulfide (CAS No. 12068-85-8)) were obtained from PubMed (https://pubmed.ncbi.nlm.nih.gov/). The bioactive compounds were screened using the following criteria in the TCMSP database: oral bioavailability (OB) ≥30% and drug-likeness (DL) value of ≥0.18. OB describes the delivery capability of orally administered drugs to the systemic circulation [14]. DL is based on the similarity in functional groups and physical properties to those of many known drugs [15]. The compounds that conformed to the OB and DL thresholds were selected as active compounds for further analyses [16].

#### 2.1.2. Screening of targets related to active compounds

Targets of active ingredients in HXZTS were obtained from the TCMSP database [13]. Since the targets provided by TCMSP may be incomplete, the targets of the active ingredients in HXZTS were predicted in Swiss Target Prediction (http://www.swisstargetprediction.ch/) [17] by using the structural information of the active ingredients obtained from PubChem (https://pubchem.ncbi.nlm.nih.gov/) (Wu et al., 2018). Finally, all obtained genes were normalized by screening in UniProt (https://www.uniprot.org/) [18].

#### 2.1.3. Collecting the therapeutic targets of OA

The therapeutic targets were attained by searching GeneCards (https://www.genecards.org/) [19] and OMIM (https://www.omim.org/) [20] with “Osteoarthritis synovium”, “osteoarthritis synovial membrane” and “osteoarthritis synovial tissue” as keywords. Then, we converted the target names to the gene names by the UniProt database [18]. In addition, the three OA synovium-related datasets (GSE55235, GSE55457, GSE12021) found in the Gene Expression Omnibus (GEO) database (https://www.ncbi.nlm.nih.gov/geo/) were merged and further the “sva” and “limma” packages (R version: 4.3.1) were used for batch correction and differentially expressed genes (DEGs) screening (|log2 (foldchange)| >1 and P value < 0.05). Using the ggplot2 and pheatmap packages, we generated volcano plots for the obtained DEGs and heatmaps for the top 50 DEGs based on expression ranking. After the removal of duplicate genes, all the OA-related targets were collected.

#### 2.1.4. Screening of common targets of HXZTS-OA

By identifying the common targets shared by OA-related targets and the HXZTS targets, a visual representation was shown in a Venn diagram using bioinformatics tools (https://www.bioinformatics.com.cn/static/others/jvenn/example.html) [21].

### 2.2 Protein-protein interaction (PPI) network construction

The construction of PPI network involved importing the common targets into the STRING database (http://string-db.org) [22]. The species was restricted to “Homo sapiens” and a confidence score of ≥0.7 was set in the interface. Cytoscape 3.9.1 (https://www.cytoscape.org/) was utilized to build a network of potential key targets and conduct a comprehensive analysis of the network parameters [23]. The CytoHubba plug-in in Cytoscape was used to identify hub genes [24].

### 2.3 Enrichment analysis of common target genes

In R software (version 4.3.1), the’clusterProfiler’ and’org.Hs.eg.db’ packages were utilized to perform Gene Ontology (GO) and Kyoto Encyclopedia of Genes and Genomes (KEGG) enrichment analyses on the obtained common target genes. The selection criterion was set to P < 0.05. The enriched terms were sorted based on their gene count, and the top 25 terms were visualized using bioinformatics tools (https://www.bioinformatics.com.cn/) [25].

### 2.4 Molecular docking

The structure files of core compounds (Quercetin, β-Sitosterol, Stigmasterol, Ginsenoside rh2, DFV, and 3α-Hydroxy-olean-12-en-24-oic-acid) were retrieved from the PubChem database (https://pubchem.ncbi.nlm.nih.gov) in SDF format, and subsequently converted to PDB files using Open Babel version 3.1.1 (http://openbabel.org/api/index.html#/api/3.0/). The 3D crystal structures of target proteins (AKT1, IL6, TNF, TP53, and VEGFA) were obtained by accessing the UniProt protein database (UniProt database: https://www.uniprot.org/). Both ligands and receptors were imported into AutoDock Vina version 1.2.2 (https://vina.scripps.edu) for dehydration, hydrogenation, and charge calculation, and then saved in pdbqt format in preparation for molecular docking. Subsequently, a docking grid was created to cover the entire target protein, and parameters were saved in txt format. Finally, molecular docking analysis was performed, and the visualization of free binding energy was presented using heat maps in bioinformatics (https://www.bioinformatics.com.cn/). Lastly, molecular docking and interactions were visualized in PyMOL.

### 2.5 Cell culture

Primary human osteoarthritic synovial fibroblasts were purchased from Procell Life Science & Technology Co., Ltd. The cells were seeded in culture dishes and cultured in a complete growth medium for human osteoarthritic synovial fibroblasts (Procell Life Science & Technology Co., Ltd, China) at 37°C with 5% CO_2_. The medium was changed every 3 days after the initial 24 h culture, and cell morphology was observed under an inverted phase-contrast microscope. Upon reaching 90% confluency, cells were passaged using 0.25% trypsin-EDTA (Gibco, USA) and used for subsequent experiments. Cells used in this study were between the 3rd and 5th passages.

### 2.6 Cell viability assay

HXZTS was purchased from Beijing Tong Ren Tang (batch number: 22101038). According to the “Chinese Pharmacopoeia” [26], the preparation method of HXZTS is as follows. The five ingredients (400 g Danggui (*Angelica sinensis* (Oliv.) Diels, DG), 80 g Sanqi (*Panax notoginseng* (Burkill) F.H.Chen, SQ), 80 g Ruxiang (*Boswellia sacra Flück.*, RX), 200 g Tubiechong (Ground Beetle, TBC), 120 g Zirantong (Pyrite, ZRT) were pulverized into the finest powder using a grinder. 20 g Bingpian (*Cinnamomum camphora* (L.) Presl, BP) was ground into a fine powder separately using a mortar and pestle. The above powders are ground, sieved, and mixed to obtain.

The cells were seeded into 6-well plates at an appropriate density and incubated overnight. After the cells adhered to the plate, they were washed twice with PBS. The medium was replaced with 1.8 ml of fresh complete growth medium and 200 µl of H_2_O (in the control group), while for the treatment group, 200 µl of HXZTS (100, 80, 60, 40, 20 mg/mL) was added. The cells were then cultured for 24 h. 10ul of CCK-8 was added into each well and incubated for 2 hours and the optical density at 450 nm was measured using a microplate reader (Varioskan flash, Thermo Scientific, USA). The experiment was replicated three times.

### 2.7 Verification using RT-qPCR

HXZTS administration was selected at a concentration that would not inhibit cell growth. After incubation, the cells were washed twice with PBS and then digested with trypsin without EDTA (Gibco, USA). The cells were centrifuged, and the pellets were collected. Total RNA extraction was performed using Trizol reagent and reverse transcription into cDNA using the PrimeScript RT Master Mix reagent kit from Takara (Japan) according to the instructions. The primer pairs used (purchased from SunYa Biotechnology Co., Ltd.) for this analysis are listed in Table 1. The GAPDH gene served as an internal control for mRNA level normalization. Reverse transcription-quantitative polymerase chain reaction (RT-qPCR) was performed using the SYBR Green Master Mix (Yeasen Bio-Technology Co., Ltd, China) and the Applied Biosystems QuantStudio 6 Flex Real-Time PCR System (Applied Biosystems, USA). The ratios of all genes’ relative mRNA expression levels were calculated with the 2^ΔΔct^ method [27].

**Table 1.**
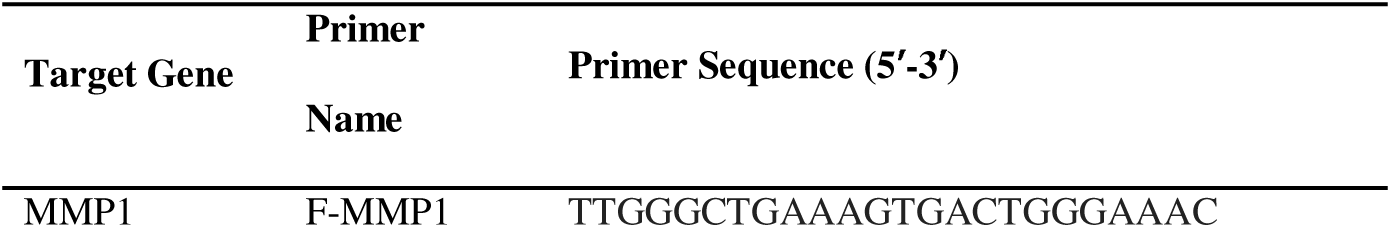
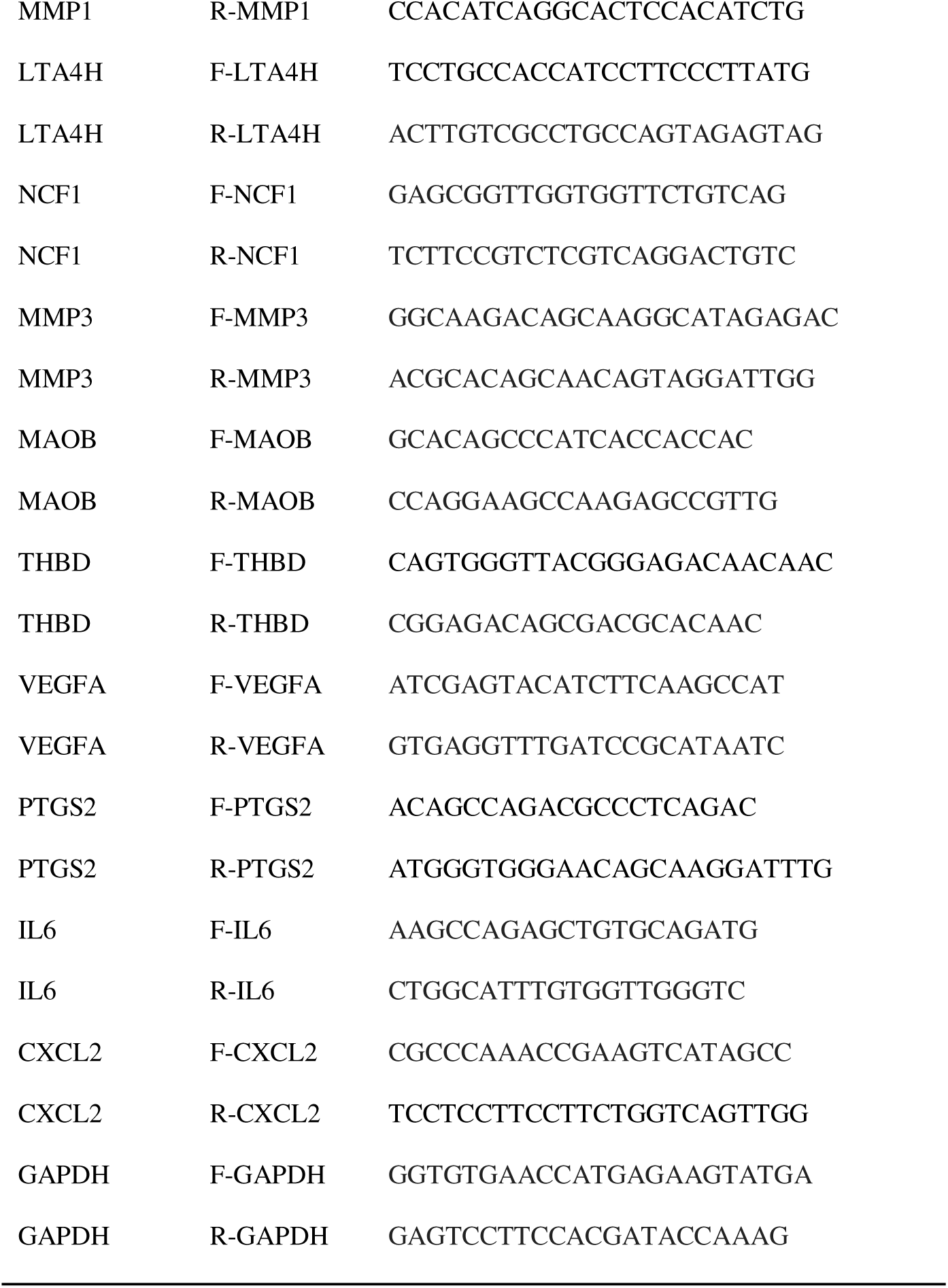
A list of primer sequences used in RT-qPCR analysis.

### 2.8 Statistical analysis

Cell activity and RT-qPCR data were presented as mean ± SD. The differences between the groups’ treatment and control were operated by unpaired t-test and were considered statistically significant at *P* < 0.05 (\**P* < 0.05, \*\**P* < 0.01, \*\*\*\**P* < 0.0001). Statistical analyses were performed using GraphPad Prism 9.0 (GraphPad Software Inc., La Jolla, CA, USA).

## 3.0 Results

### 3.1 Potential targets of active ingredients

Utilizing the criteria of OB ≥ 30% and DL ≥ 0.18, 21 active ingredients in HXZTS were identified through the TCMSP database. These include two ingredients from DG, eight from SQ, eight from RX, and three from BP. Venn diagrams showed the numbers of active compounds in each herb of HXZTS (Fig. 2A). Comprehensive details of these active ingredients can be found in Supplementary Table S1. Using the TCMSP target module, 170 potential targets of HXZTS active compounds and their corresponding gene symbols were identified, as detailed in Supplementary Table S2.

**Fig. 2.**
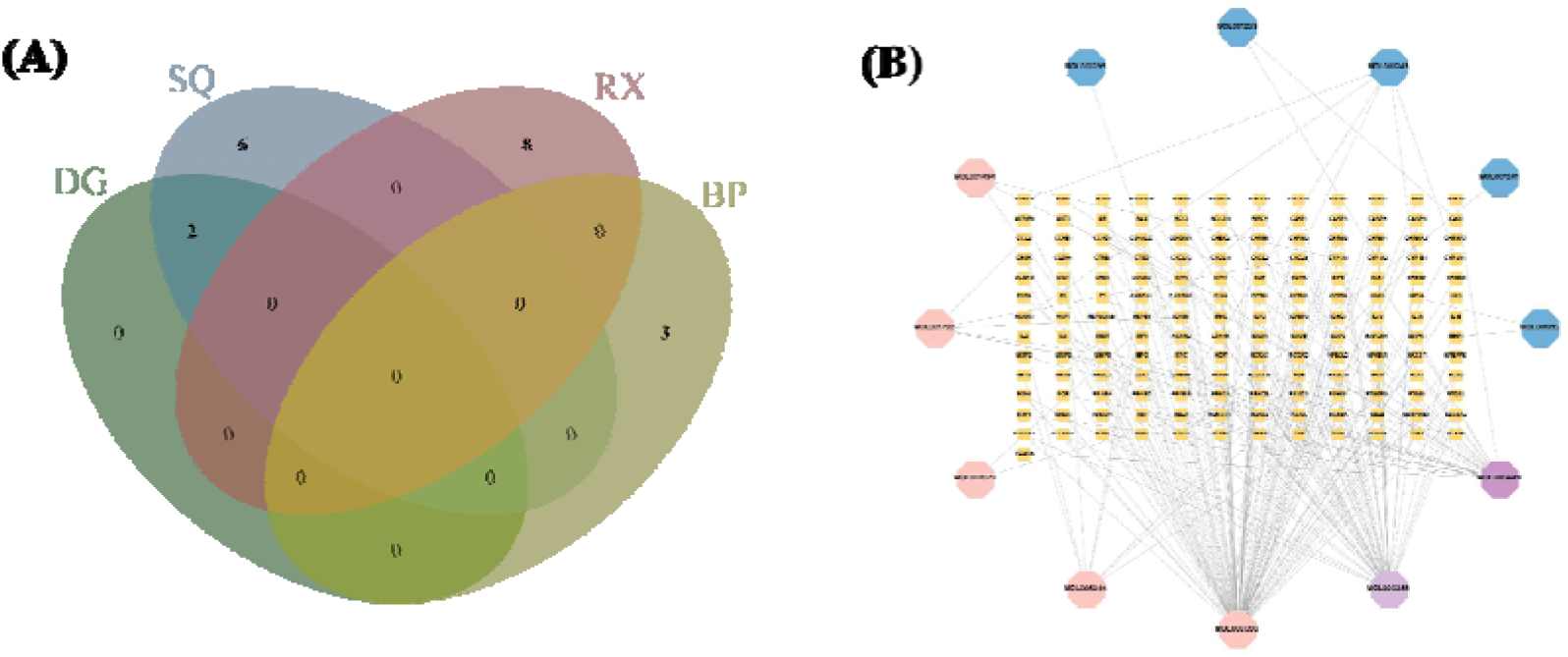
Screening for active compounds and targets of HXZTS. (A) Venn diagram shows the number of active compounds in each of the HXZTS herbs. In green is DG (2 compounds), in blue is SQ (8 compounds), in red is RX (8 compounds), and in yellow is BP (3 compounds). (B) Compound-target network. Octagons represent compounds in HXZTS, blue octagonal nodes are RX compounds, red octagonal nodes are SQ compounds, purple octagonal nodes are DG compounds, and yellow square nodes represent predicted targets. The edges represent the interaction between the compound and the target, and the node size is proportional to the degree of interaction.

### 3.2 Compound-target network

These compounds and targets were integrated into Cytoscape 3.9.1 to construct an herb–compound– target network, which consists of 169 nodes and 207 edges (Fig. 2B). This network elucidated the multifaceted effects of HXZTS’s active compounds on OA. Network analysis revealed that several compounds, including Quercetin (MOL000098), β-Sitosterol (MOL000358), Stigmasterol (MOL000449), Ginsenoside rh2 (MOL005344), DFV (MOL001792), and 3α-Hydroxy-olean-12-en-24-oic-acid (MOL001243), are key nodes.

### 3.3 Common targets of HXZTS-OA

In our study, we searched for differentially expressed genes (DEGs) in osteoarthritis (OA) by comparing the levels of gene expression between OA patients and normal synovium samples in our test group. Through the analysis of three series of data (GSE12021, GSE55235, and GSE55457), we identified a total of 441 genes, with 224 genes found to be up-regulated and 217 genes found to be down-regulated in OA (Supplementary Table S3). The expression patterns of the top 50 DEGs out of 441 are visualized in Fig. 3B using heatmaps. Additionally, we integrated disease targets obtained from GeneCards, OMIM, and the GEO DEGs, removing any duplicates, resulting in a total of 2062 disease targets specific to OA (Supplementary Table S4). By identifying the intersection between the active compound targets of HXZTS and the disease targets of OA, we identified 111 common targets. These common targets were selected as key targets for evaluating the anti-OA activity of the HXZTS compounds, as illustrated in Fig. 3C and described in Supplementary Table S5.

**Fig. 3.**
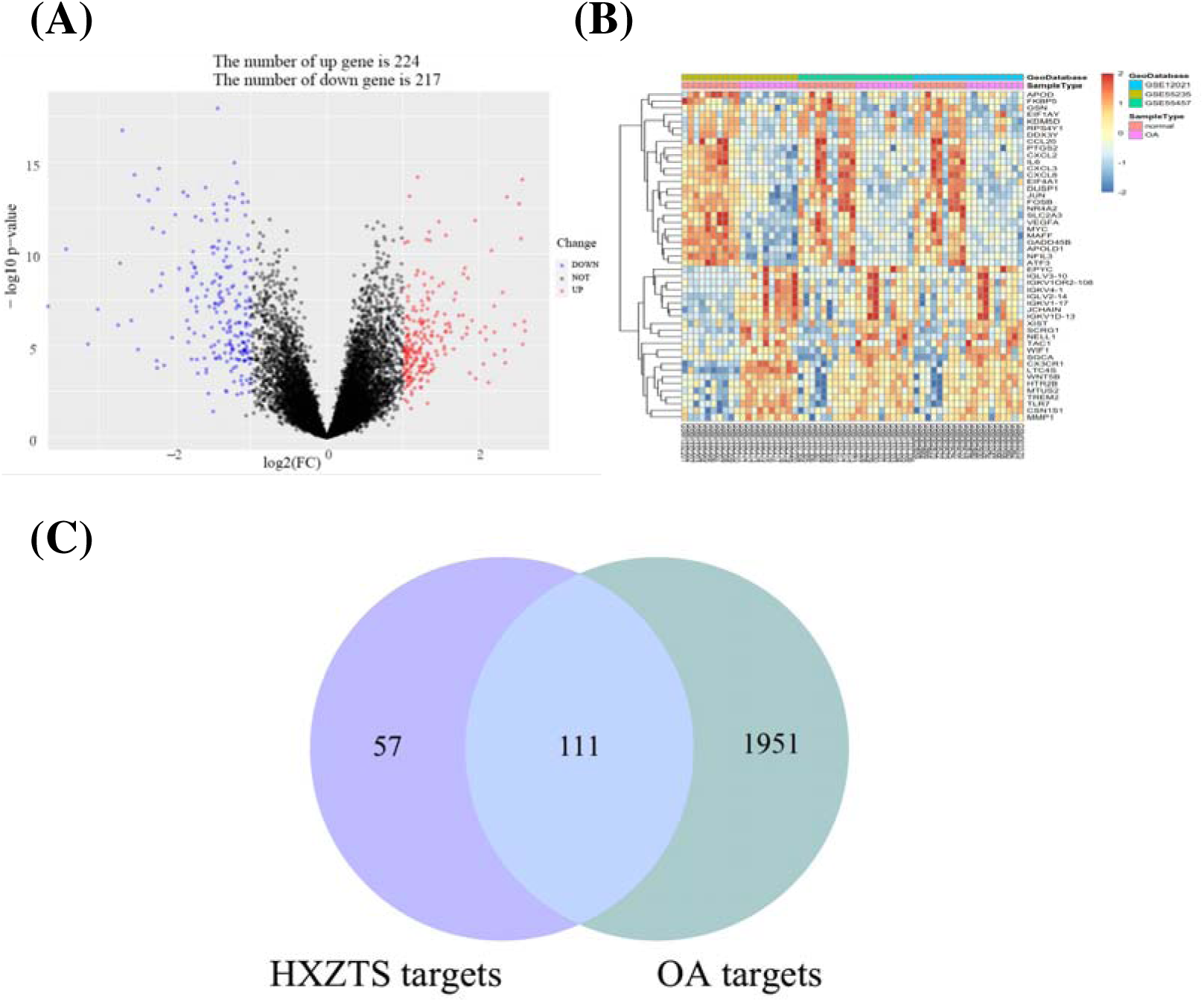
Screening for common targets of HXZTS-OA. (A) The differential gene volcano plot shows the distribution of genes in synovium samples. Red and green represent up-regulated and down-regulated genes, respectively, while black indicates no significant differences. (B) Heat map showing the expression patterns of the top 50 DEGs out of 441. Columns correspond to samples and rows correspond to genes. Each batch represents a GEO series. Batch 1: GSE12021; Batch 2: GSE55235; Batch 3: GSE55457. (C) Venn diagram of 111 common targets between HXZTS active compound targets and OA disease targets.

### 3.4 PPI network of common targets

To explore the mechanism of HXZTS in treating OA, we imported 111 common targets into the STRING database to construct a protein-protein interaction (PPI) network. This network includes 110 nodes and 2090 edges, with a degree centrality (DC) of 72. We identified the core PPI network based on topological analysis, using a ≥2-fold median DC as the filtering criterion. Through network analysis, we identified 11 core targets (Fig. 4 (left)). We constructed networks of core and non-core targets, where node size and color intensity are proportional to the target degree. Using the CytoHubba plugin, we selected the top 10 genes ranked by the MCC method as hub genes, which are AKT1, TNF, TP53, IL6, CASP3, PTGS2, IL1B, HIF1A, VEGFA, EGFR. (Fig. 4 (right), Supplementary Table S6).

**Fig. 4.**
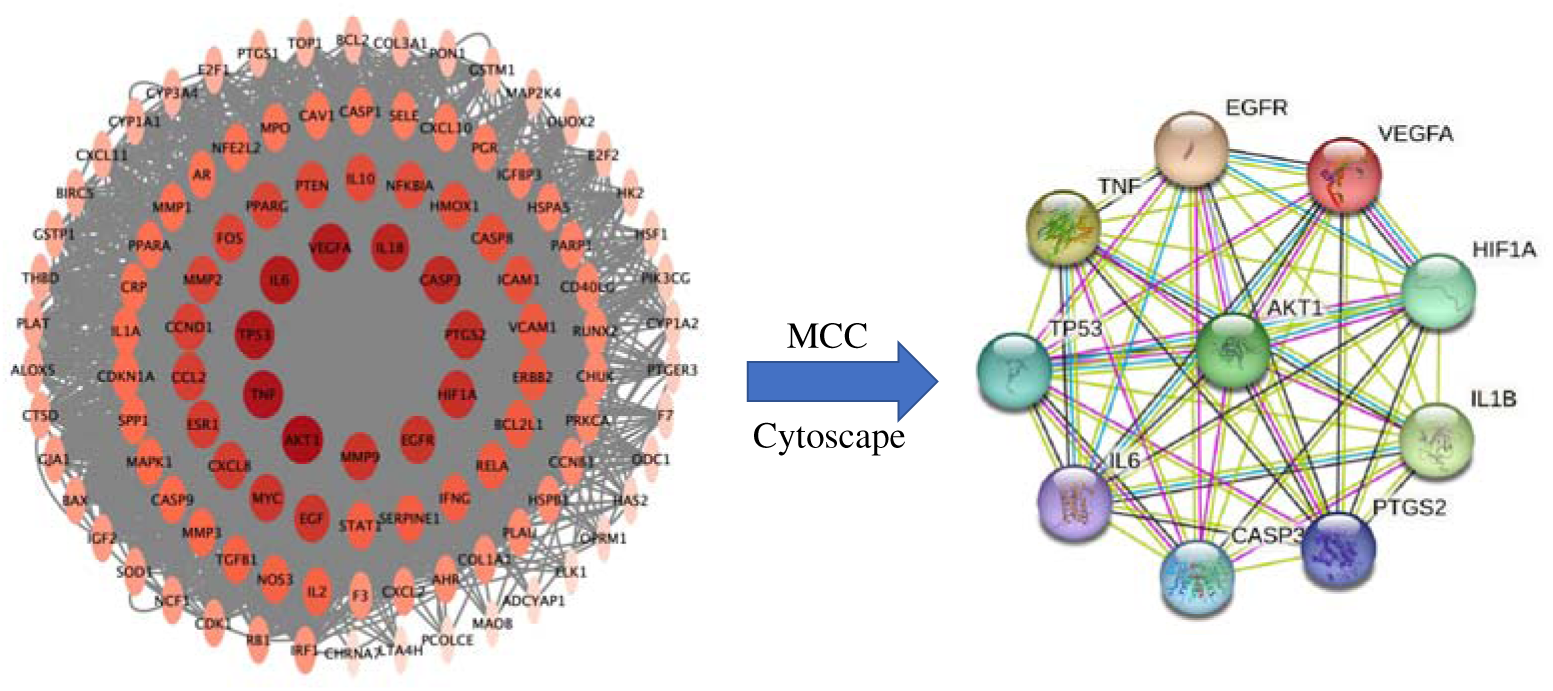
Identification of hub genes of HXZTS for OA. PPI network of 111 common targets. This network has 111 nodes and 2090 edges. Node size and color are proportional to the degree of interaction (left). PPI network of the hub genes using the CytoHubba (right).

### 3.5 GO enrichment analysis of common targets

To further explore the various mechanisms of HXZTS in treating OA, we conducted Gene Ontology (GO) enrichment analysis on the 110 common targets. A total of 2126 terms were identified (Supplementary Table S7-9), including 1947 biological processes (BP), 131 molecular functions (MF), and 48 cellular components (CC). As shown in Fig. 5A-C, the top 25 enriched BP, MF, and CC terms are presented in bubble charts. The enriched BP terms primarily involve response to molecule of bacterial origin (GO:0002237), response to lipopolysaccharide (GO:0032496), response to oxidative stress (GO:0006979), regulation of apoptotic signaling pathway (GO:2001233), regulation of inflammatory response (GO:0050727), etc. Highly enriched MF terms include DNA-binding transcription factor binding (GO:0140297), cytokine receptor binding (GO:0005126), ubiquitin-like protein ligase binding (GO:0044389), endopeptidase activity (GO:0004175), etc. Additionally, CC terms include membrane raft (GO:0045121), membrane microdomain (GO:0098857), vesicle lumen (GO:0031983), collagen-containing extracellular matrix (GO:0062023), secretory granule lumen (GO:0034774), etc.

**Fig. 5.**
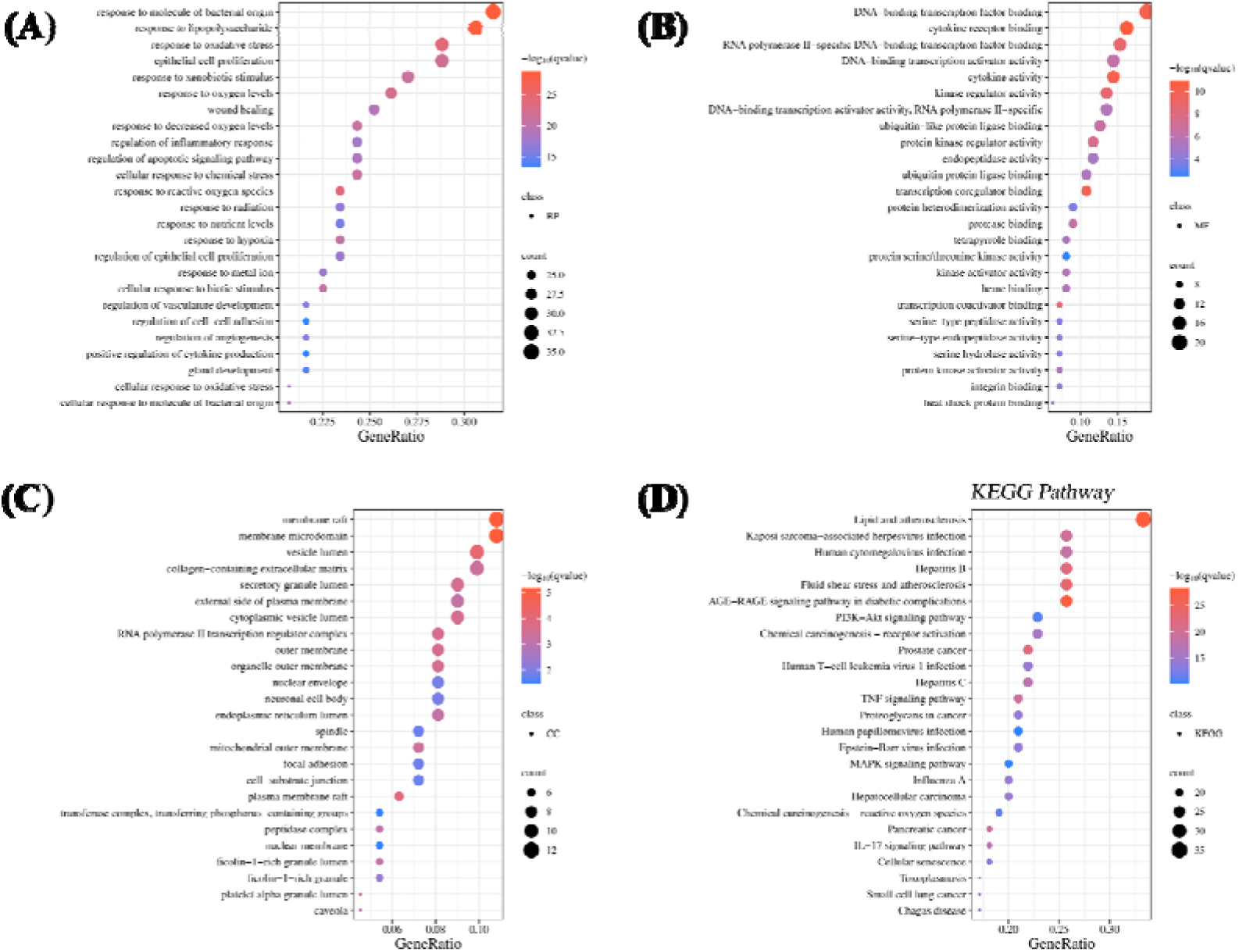
GO enrichment analysis of the therapeutic target. (A) BP enrichment analysis of common targets (the top 25 results). (B) MF enrichment analysis of common targets (the top 25 results). (C) CC enrichment analysis of common targets (the top 25 results). (D) KEGG enrichment analysis of common targets (the top 25 results).

A KEGG pathway enrichment analysis was conducted on 111 common targets, applying a significance level of q-value < 0.05. This analysis revealed that the targets were significantly enriched in 154 pathways (as detailed in Supplementary Table S10). The top 25 most highly enriched pathways were then selected based on the number of associated genes, as shown in Fig. 5D. The results of the KEGG analysis revealed the involvement of several critical pathways, underscoring the multifunctional nature of the observed effects. Specifically, the analysis identified anti-inflammatory responses through the TNF signaling pathway (hsa04668), IL-17 signaling pathway (hsa04657), and MAPK signaling pathway (hsa04010), indicating a broad spectrum of anti-inflammatory activity [28–30]. Additionally, cell protection was highlighted through the PI3K-Akt signaling pathway (hsa04151), suggesting a mechanism for cellular defense and survival [31]. The analysis also pointed to anti-infection capabilities associated with Hepatitis B (hsa05161), Hepatitis C (hsa05160), and Human T-cell leukemia virus 1 infection (hsa05166), reflecting potential antiviral effects [32, 33]. Furthermore, metabolic regulation, particularly in lipid metabolism and atherosclerosis (hsa05417), was identified, indicating a role in modulating metabolic processes [34]. Finally, the mitigation of oxidative stress was observed in the context of chemical carcinogenesis and the generation of reactive oxygen species (hsa05207), highlighting antioxidant properties [35]. These pathways collectively illustrated a comprehensive therapeutic potential, addressing inflammation, infection, metabolic health, and oxidative stress.

### 3.6 Verification and visualization of the molecular docking results

Based on the results from PPI analysis, six compounds were selected for their high degree values as core active components: Quercetin (MOL000098), β-Sitosterol (MOL000358), Stigmasterol (MOL000449), Ginsenoside rh2 (MOL005344), DFV (MOL001792), and 3α-Hydroxy-olean-12-en-24-oic-acid (MOL001243). These compounds were subjected to molecular docking with five core targets identified by CytoHubba: AKT1, TNF, TP53, IL6, and VEGFA. The molecular docking results (Fig. 6A and Supplementary Table S11) showed that the binding energies between the core active components and the core targets were all less than-5.0 kcal/mol, indicating good binding activity. Notably, the interactions between IL6 and Quercetin; TNF and 3α-Hydroxy-olean-12-en-24-oic-acid; AKT1, TP53, and all six compounds demonstrated binding energies of less than-7.0 kcal/mol, suggesting a strong binding affinity.

**Fig. 6.**
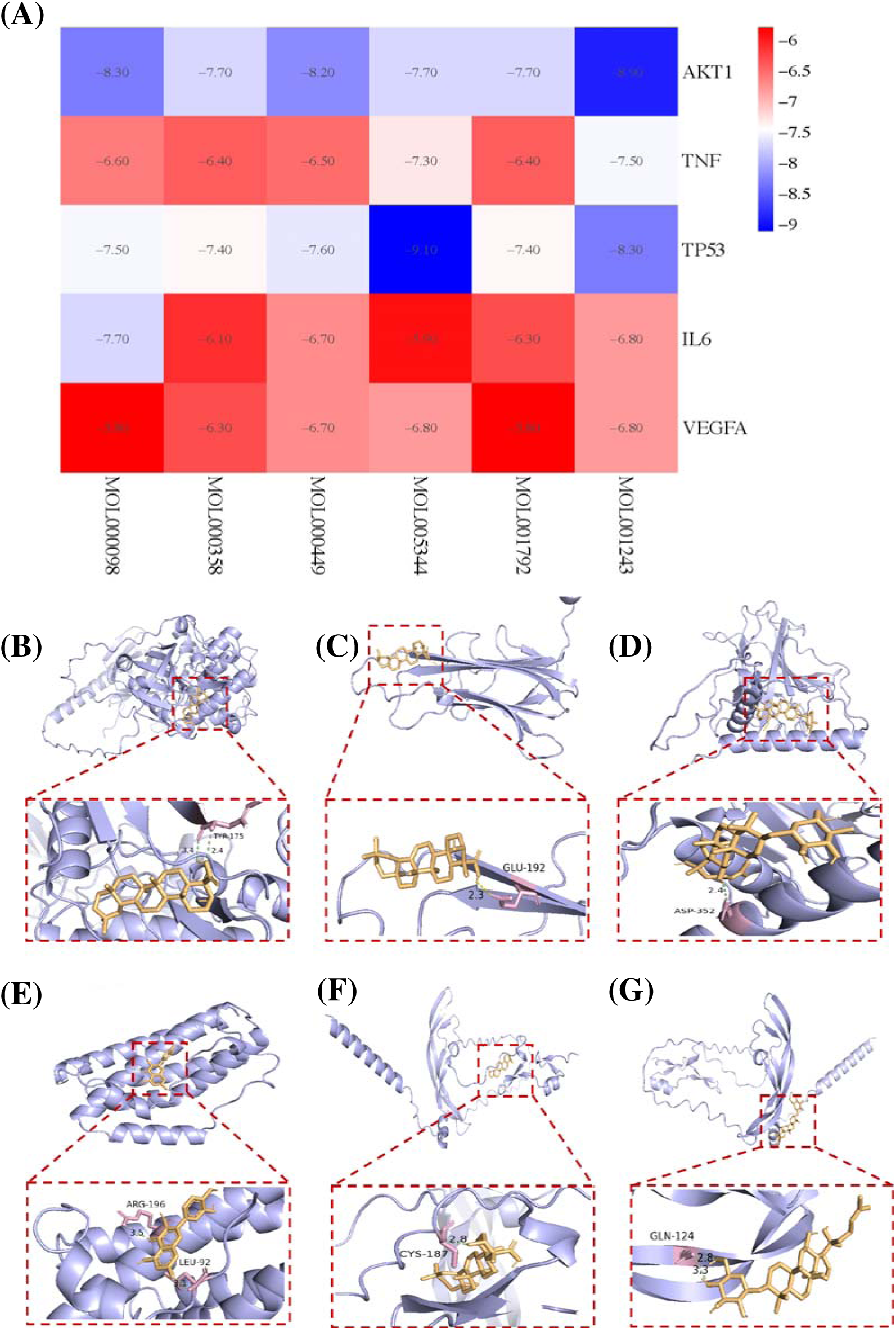
Molecular docking of key targets and specific active compounds. (A) Heat map of molecular docking score (Binding energy(kcal/mol) of key targets and active compounds of herbs); Docking patterns of key targets and specific active compounds. 3α-Hydroxy-olean-12-en-24-oic-acid-AKT1 (B), 3α-Hydroxy-olean-12-en-24-oic-acid-TNF (C), Ginsenoside rh2-TP53 (D), Quercetin-IL6 (E), 3α-Hydroxy-olean-12-en-24-oic-acid-VEGFA (F), and Ginsenoside rh2-VEGFA (G).

For visualization, the docking results with the greatest binding energy between the core active components and targets were selected, and diagrams were generated using PyMOL software (Fig. 6B-G). As illustrated, AKT1 can dock to 3α-Hydroxy-olean-12-en-24-oic-acid through hydrogen bond interaction with the amino acid residue TYR-175. TNF can dock to 3α-Hydroxy-olean-12-en-24-oic-acid through a hydrogen bond with the amino acid residue GLU-192. TP53 can dock to Ginsenoside rh2 through a hydrogen bond with the amino acid residue ASP-352. IL6 can dock to Quercetin through hydrogen bonds with amino acid residues ARG-196 and LEU-92. VEGFA can dock to 3α-Hydroxy-olean-12-en-24-oic-acid and Ginsenoside rh2 through hydrogen bonds with amino acid residues CYS-187 and GLN-124, respectively.

### 3.7 Cell viability after HXZTS treatment

To determine human osteoarthritic synovial fibroblast viability under the treatment of HXZTS, a CCK-8 cell viability assay was performed. Cells were treated with HXZTS that had undergone a gradient dilution of 20∼100 mg/ml for 24h. Our data showed that 40-100mg/ml HXZTS has a certain inhibitory effect on the viability of synovial fibroblasts (Fig. S1). However, 20 mg/ml HXZTS slightly promoted the viability of synovial fibroblasts. Therefore, HXZTS with a 20 mg/ml concentration was selected for subsequent experiments.

### 3.8 Verification by RT-qPCR

To validate the therapeutic targets of HXZTS identified through network pharmacology and bioinformatics analysis for the treatment of synovitis in osteoarthritis, RT-qPCR was utilized to assess the mRNA expression levels in human osteoarthritic synovial fibroblasts after HXZTS treatment.

Network pharmacology and bioinformatics analysis identified 110 common targets for HXZTS in the treatment of synovitis in osteoarthritis. After the intersection with the 440 targets obtained from GEO data on synovitis in osteoarthritis, 20 common targets were obtained (Fig. S2 and Supplementary Table S12). Subsequently, RT-qPCR validation of five randomly selected upregulated genes and five randomly selected downregulated genes revealed significant alterations in mRNA expression levels post-treatment, with normalization against GAPDH for accurate quantification. Our RT-PCR results showed that the expression level of MMP1, LTA4H, NCF1, MMP3, and MAOB was upregulated, while the expression levels of THBD, VEGFA, PTGS2, IL6, and CXCL2 were downregulated (Fig. 7A). Specifically, when compared to the control group, the treatment group exhibited significant differences in the RNA expression levels of MMP1, LTA4H, NCF1, MAOB, PTGS2, IL6, and CXCL2. The RT-qPCR and GEO data exhibited a high degree of concordance (*R^2^*=0:8175), affirming the reliability of the RT-qPCR findings and the potential biological relevance of these genes in response to HXZTS treatment in osteoarthritic synovitis (Fig. 7B).

**Fig. 7.**
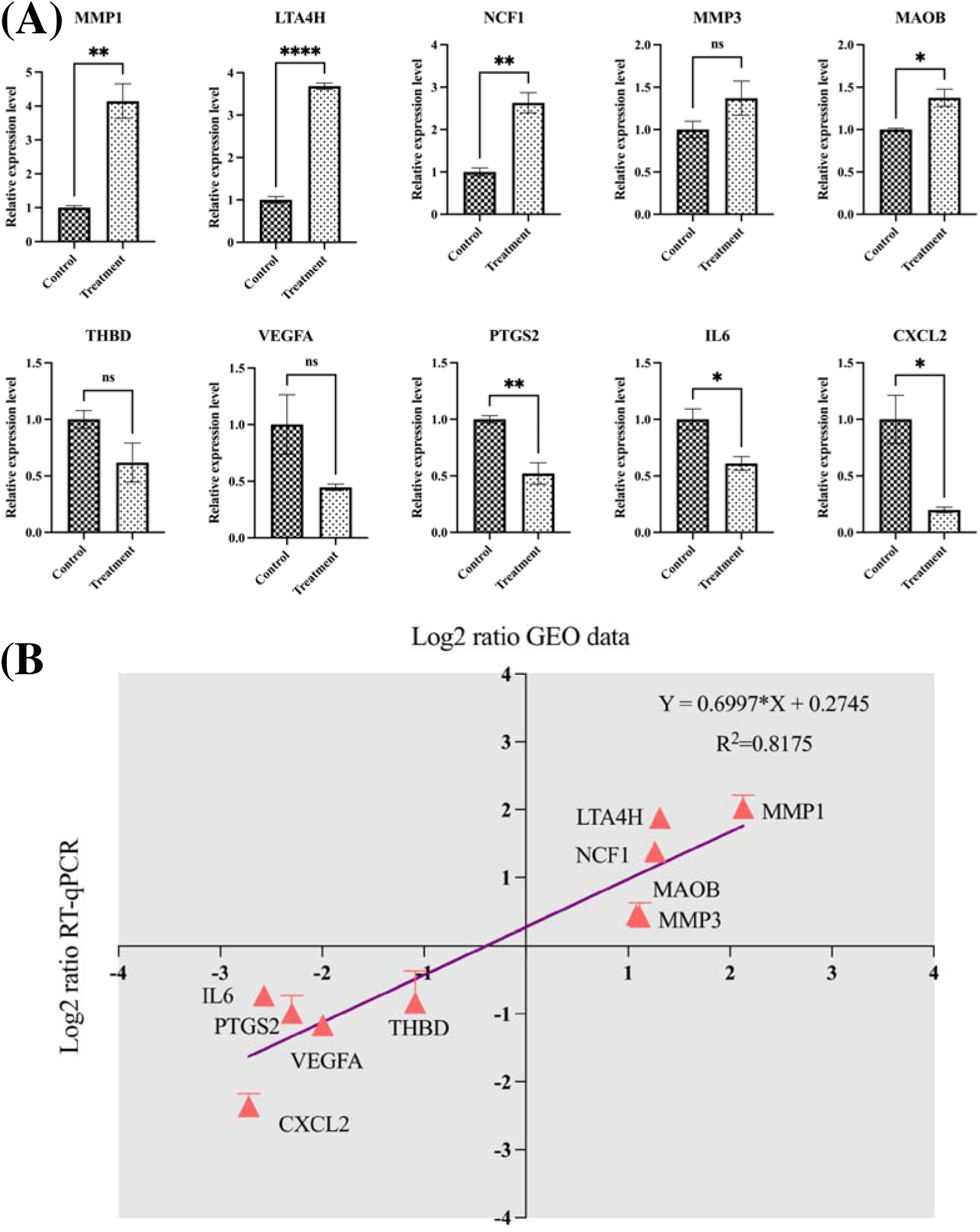
**Differential target mRNA expression and correlation analysis in human osteoarthritic synovial fibroblasts post-HXZTS treatment. (**A) mRNA expression levels in human osteoarthritic synovial fibroblasts after HXZTS treatment by RT-qPCR normalized to GAPDH. Data represent mean ± SD (n = 3; \**P* < 0.05, \*\**P* < 0.01, \*\*\*\**P* < 0.0001, t-test). (B) Correlation between GEO data and RT-qPCR data. The X-axis of the plot represents the log_2_(Fold Change) of GEO data, while the y-axis represents the log_2_(Fold Change) of RT-PCR data. The *R^2^* value refers to the coefficient of determination of the regression line.

## 4. Discussion

OA is a degenerative joint disease influenced by multifactorial and intricate interactions [36]. Currently, there remains a dearth of widely accepted and clinically applicable biomarkers for the diagnosis of OA. While previous research has predominantly focused on the degenerative changes in articular cartilage in OA, emerging evidence underscores the pivotal roles of synovial inflammation and dysfunction of synovial fibroblasts in the pathogenesis of this disease. Elucidating the molecular mechanisms underlying synovial tissue damage in osteoarthritis is crucial for the development of targeted therapeutic interventions capable of effectively modulating disease progression and alleviating symptoms [37]. HXZTS, a traditional Chinese medicine formula, has shown promising therapeutic effects in the treatment of osteoarthritis, as observed by previous studies [8]. However, its molecular mechanisms of action, particularly on the synovial tissue, remained unclear until now. Our study integrates network pharmacology, bioinformatics analysis, molecular docking, and experimental validation to provide valuable insights into the potential targets, pathways, and mechanisms by which HXZTS exerts therapeutic effects on synovial tissue in osteoarthritis.

The identification of 21 active ingredients from HXZTS, with specified origin from DG, SQ, RX, and BP, and their subsequent assembly into a compound-target network, establishes a foundation for understanding the multi-target interactions of HXZTS in OA treatment. This network, comprising 169 nodes and 207 edges, is demonstrative of the complex nature of the pharmacological effects of HXZTS, with several key compounds, such as Quercetin, β-Sitosterol, Stigmasterol, serving as central nodes within the network. The elucidation of 111 common targets between HXZTS and OA by intersecting the active compound targets with disease-specific targets indicates the therapeutic potential of HXZTS on osteoarthritic synovial tissue [38]. These common targets form a robust scaffold for evaluating the therapeutic effects of HXZTS on osteoarthritic synovial tissue, which is further substantiated by the construction of a comprehensive PPI network and subsequent identification of 11 core targets. This finding clearly shows the potential therapeutic efficacy of HXZTS in treating the pathological processes of osteoarthritic synovial tissue, providing a strong foundation for further elucidating its underlying mechanisms of action.

Moreover, the GO enrichment analysis of these targets revealed significant biological processes like response to oxidative stress, regulation of inflammatory response, apoptosis regulation, and cytokine signaling. Notably, the enrichment analysis indicated that HXZTS could alleviate inflammation in osteoarthritic synovial tissue through the TNF signaling pathway, IL-17 signaling pathway, and MAPK signaling pathway. These pathways are critically involved in the pathogenesis of synovial inflammation and joint destruction in OA [39, 40]. The molecular docking results further demonstrated the strong binding affinities of HXZTS active compounds, such as quercetin and 3α-hydroxy-olean-12-en-24-oic acid, with key inflammatory targets like TNF, IL6, and AKT1. Notably, the RT-qPCR validation confirmed the downregulation of IL6 and PTGS2 as well as the upregulation of MMP1 and MMP3 upon HXZTS treatment, corroborating its anti-inflammatory and potential cartilage repair effects [41–44]. The upregulation of LTA4H and NCF1, involved in oxidative stress and inflammation, suggests HXZTS may activate certain antioxidant pathways. In addition, the enrichment of oxidative stress-related pathways, such as the generation of reactive oxygen species and chemical carcinogenesis, suggests that HXZTS may exert antioxidant effects on osteoarthritic synovial tissue. Oxidative stress has been implicated in the pathogenesis of OA, contributing to cartilage degradation, synovial inflammation, and subchondral bone remodeling [45]. HXZTS also might modulate apoptosis in osteoarthritic synovial tissue, as evidenced by the enrichment of the apoptotic signaling pathway and the strong binding of HXZTS compounds with the apoptosis-related target TP53. Aberrant apoptosis of synovial fibroblasts has been linked to synovial hyperplasia and joint inflammation in OA [46]. The downregulation of VEGFA upon HXZTS treatment, as validated by RT-qPCR, may also contribute to the modulation of angiogenesis and synovial hyperplasia [47]. Furthermore, the enrichment of pathways related to metabolic regulation, such as lipid metabolism and atherosclerosis, suggests that HXZTS may also exert beneficial effects on metabolic dysfunction associated with OA. Metabolic alterations, including lipid dysregulation, have been implicated in the pathogenesis of OA and may contribute to synovial inflammation and joint degeneration [48].

Collectively, these findings suggest that HXZTS exerts its therapeutic effects on osteoarthritic synovial tissue through a multifaceted mechanism involving the modulation of inflammation, oxidative stress, apoptosis, angiogenesis, and metabolic regulation. The active compounds of HXZTS, such as quercetin, β-sitosterol, stigmasterol, ginsenoside rh2, DFV, and 3α-hydroxy-olean-12-en-24-oic acid, interact with key targets like AKT1, TNF, TP53, IL6, and VEGFA, thereby influencing various signaling pathways implicated in OA pathogenesis. From a clinical perspective, these findings provide insights into the potential application of HXZTS as a complementary or alternative therapeutic approach for managing osteoarthritis, particularly in alleviating synovial inflammation and associated symptoms. By targeting multiple pathways and processes involved in OA pathogenesis, HXZTS may offer a more comprehensive and holistic treatment strategy compared to conventional therapies that primarily focus on symptom management. Furthermore, the identified mechanisms and targets could serve as a foundation for the development of novel targeted therapies or the optimization of existing treatments for OA. It is important to note that while this study provides valuable insights into the potential mechanisms of HXZTS in osteoarthritic synovial tissue, further in vivo studies and clinical trials are warranted to validate these findings and evaluate the efficacy and safety of HXZTS in the treatment of osteoarthritis. Additionally, explorations into the synergistic effects of the various active compounds and their interactions with multiple targets could further enhance our understanding of the therapeutic potential of HXZTS.

## 5. Conclusion

Our study elucidates the potential mechanisms of HXZTS in modulating osteoarthritic synovial tissue through the integration of network pharmacology, bioinformatics analysis, molecular docking, and experimental validation. The findings suggest that HXZTS exerts its therapeutic effects by modulating multiple pathways and targets associated with inflammation, oxidative stress, apoptosis, angiogenesis, and metabolic regulation. The results obtained through the bioinformatics and computational analyses provide compelling evidence that quercetin, β-sitosterol, stigmasterol, ginsenoside rh2, DFV, and 3α-hydroxy-olean-12-en-24-oic acid may constitute the principal bioactive ingredients of HXZTS responsible for its therapeutic efficacy on OA synovium. These active compounds interact with key targets such as AKT1, TNF, TP53, IL6, and VEGFA, thereby influencing various signaling pathways implicated in the pathogenesis of osteoarthritis. These insights contribute to our understanding of the pharmacological basis of HXZTS in treating osteoarthritis and provide a foundation for developing novel targeted therapies and optimizing existing treatment strategies for this debilitating condition.

## Glossary

OA: Osteoarthritis
TCM: Traditional Chinese Medicine
HXZTS: Huoxuezhitongsan
DG: Danggui (Angelica sinensis (Oliv.) Diels)
SQ: Sanqi (Panax notoginseng (Burkill) F.H.Chen)
RX: Ruxiang (Boswellia sacra Flück.)
BP: Bingpian (Cinnamomum camphora (L.) Presl)
TBC: Tubiechong (Ground Beetle)
ZRT: Zirantong (Pyrite)
OB: Oral bioavailability
DL: Drug-likeness
GEO: Gene Expression Omnibus database
DEGs: Differentially expressed genes
PPI: Protein–protein Interaction
GO: Gene Ontology
KEGG: Kyoto Encyclopedia of Genes and Genomes
BP: Biological Process
CC: Cellular Component
MF: Molecular functions
RT-qPCR: Reverse transcription-quantitative polymerase chain reaction

## Acknowledgments

This study was financially supported by the Natural Science Foundation of Fujian Province (Nos. 2022J011310, 2023J011542, 2023J011547); Fujian Provincial Clinical Medical Research Center for First Aid and Rehabilitation in Orthopaedic Trauma (Nos. 2020Y2014); 2022 Fuzhou Science and Technology Plan technology innovation platform project (Nos. 2022-P-018); 2021 Fuzhou health innovation platform construction project (Nos. 2021-S-wp2).

## Authors’ contributions

**Li Chen:** Conceptualization, Methodology, Formal analysis, Investigation, Writing - Original Draft. **Hongxiu Wang:** Investigation, Visualization, Funding acquisition. **Dongzhi Wu:** Investigation, Resources. **Jinlan Su, Shunxi Chen:** Visualization, Writing - Review & Editing. **Dongdong Chen:** Writing - Review & Editing, Funding acquisition. **Tao Zhang:** Writing - Review & Editing, Funding acquisition. **Wenhui He:** Conceptualization, Writing - Review & Editing, Funding acquisition

## Additional information

Please refer to Supplementary Figures and Supplementary Tables.

