## Supplemental Figure for "Network pharmacology, bioinformatics, molecular docking and experimental verification of the mechanism of HXZTS treatment on osteoarthritis synovium"

Title:

Integrating network pharmacology, molecular docking, and experimental validation for interpreting the multitarget mechanisms of HXZTS in osteoarthritic synovial tissue

Authors:

Li Chen ^1,2^†, Hongxiu Wang^1,3^†, Dongzhi Wu^1^, Jinlan Su^1,3^, Shunxi Chen^1,3^, Dongdong Chen^1,4^, Hong Zheng^1,4^, Tao Zhang^1^*, and Wenhui He^1^*

Affiliations:

^1^Department of Orthopaedics Institute, Fuzhou Second General Hospital, Fuzhou, Fujian 350007, People’s Republic of China

^2^Institute of Biological Sciences, Faculty of Science, Universiti Malaya, 50603 Kuala Lumpur, Malaysia.

^3^Department of Rehabilitation Medicine, Fuzhou Second General Hospital, Fuzhou, Fujian 350007, People’s Republic of China

^4^Fujian Provincial Clinical Medical Research Center for First Aid and Rehabilitation in Orthopaedic Trauma, Fuzhou, Fujian 350007, People’s Republic of China

† These authors contributed equally to this work and share the first authorship

* Correspondence:

 (Tao Zhang); (Wenhui He)

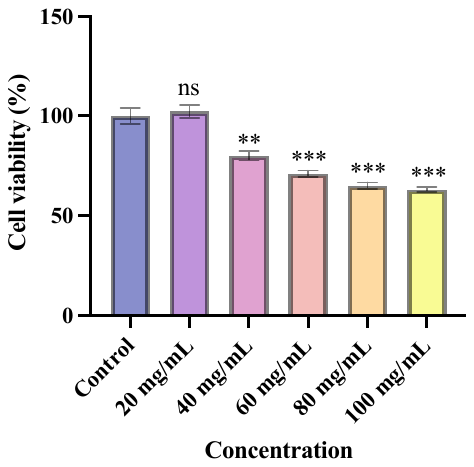

**Fig. S1. Cell viability treated with HXZTS at different concentrations.**

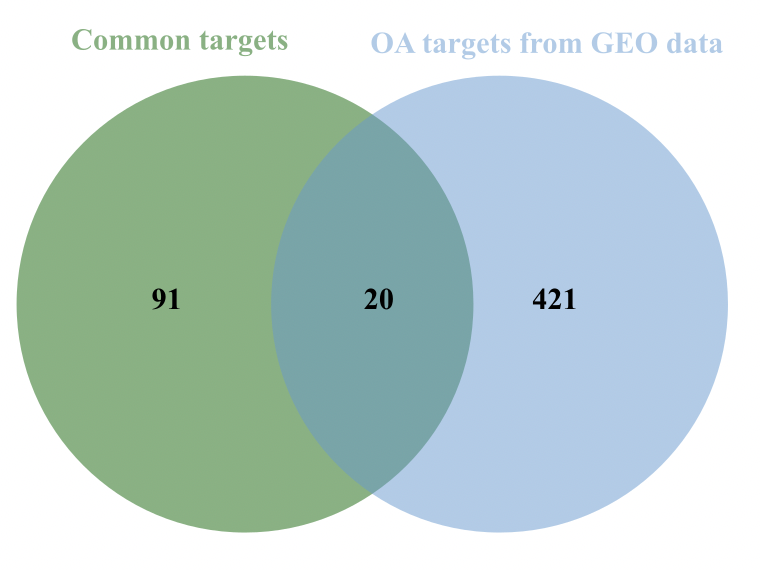

**Fig. S2. Venn diagram of common targets and OA targets from GEO data.**
